# Chili Pepper and Synthetic Capsaicinoid Flavourants in “Heat” and “Unflavoured” Oral Nicotine Pouches Marketed in the United States

**DOI:** 10.64898/2026.08.19.745863

**Authors:** Sairam V. Jabba, Zhong Li, Sven E. Jordt

**Author notes:** Corresponding Author: Sven E. Jordt, PhD, Department of Anesthesiology, Duke University School of Medicine, 3 Genome Ct., Durham, NC 27710-3094, USA.

## Abstract

**Background:** In the United States, several states have restricted sales of flavoured tobacco products, including popular menthol-and mint-flavoured Oral Nicotine Pouches (ONP). In response, tobacco companies introduced “unflavoured” ONP containing odorless synthetic cooling agents. Since these, in turn, have become targets of legislative bans, the tobacco industry may seek out flavourants with other sensory effects to increase the appeal of “unflavoured” ONP.

**Methods:** Online merchants were searched for “unflavored” ONP marketed to consumers in jurisdictions with flavour bans. Sensory effects of aqueous extracts from identified “heat”, “spicy” and “unflavoured” ONP were analyzed by Ca^2+^-microfluorimetry in HEK293 cells expressing the human heat/chili pepper flavourant (capsaicinoid) receptor, hTRPV1. ONP were analyzed for capsaicinoids and sweeteners by Liquid-Chromatography/Mass-Spectrometry (LC/MS).

**Results:** A new category of “heat” or “spicy” ONP was identified, including products marketed as “unflavoured”. Extracts from all these ONP robustly activated TRPV1, with “unflavoured” Lucy Heat the most potent. Chemical analysis demonstrated that Lucy Heat contained the synthetic capsaicinoid nonivamide at high levels (∼675 µg/pouch), while others contained mixtures of capsaicinoids (5-25 µg/pouch) combined with other characterizing flavours (tropical, fruit). *ONP tested for sweeteners contained either sucralose or acesulfame-k*.

**Conclusions:** The tobacco industry continues to probe regulatory loopholes by claiming that newly introduced capsaicinoids and sweeteners in ONP do not represent characterizing flavours. This is contradicted by industry and regulatory determinations assigning characterizing flavour properties to these additives. The toxicological health risks of repeated capsaicinoid exposures due to ONP use, in combination with nicotine and other constituents, need to be assessed.

- **What is already known on this topic** – Tobacco companies employ chemical strategies to overcome regulatory restrictions on flavours introduced to protect youth, exemplified by the replacement of menthol and mint flavourants in Oral Nicotine Pouches (ONP) with synthetic cooling agents.
- **What this study adds** – This study reveals a new industry strategy to bypass flavour bans in the United States, introducing chili pepper flavourants to increase the appeal of “unflavoured” ONP marketed in jurisdictions in which cooling agents are banned.
- **How this study might affect research, practice or policy** – The regulatory status of hot chili pepper flavourants in tobacco products needs to be clarified and their potential toxicity investigated, especially in combination with nicotine.

## INTRODUCTION

Sales of oral nicotine pouches (ONP) have increased dramatically in recent years in the United States and world-wide. ^1–3^ ONP use has been rising across all age groups, particularly among youth and young adults.^4–6^ In the United States, mint and menthol are the most popular ONP flavour varieties among adolescents and adults.^7–9^ Mint and menthol flavours are imparted by flavour chemicals such as menthol, menthone and carvone, originally derived from mint plants such as peppermint or spearmint. These flavourants create minty odor sensations through the olfactory sense and elicit cooling sensations through a mechanism named chemesthesis, the activation of physical sensations by chemical stimuli.^10^ ^11^ Specifically, mint flavourants trigger the sensation of cooling by activating the cold/menthol receptor Transient Receptor Potential Melastatin 8 (TRPM8) on peripheral sensory nerves that transmit changes in internal and external temperature to the central nervous system.^12^ The chemesthetic cooling properties of menthol and mint chemicals significantly contribute towards the appeal, initiation and consumption of tobacco products.^13^

The US federal states California and Massachusetts and the District of Columbia (DC) banned tobacco products with characterising flavours, including mint and menthol, with the exemption of tobacco flavour. These policies also apply to ONP.^14^ ^15^ Tobacco companies have challenged state flavour bans by introducing ONP advertised as “unflavoured” or “flavour ban-approved”. However, some of these ONP still contain appealing flavourants, including sweeteners and chemesthetic chemicals.^16^ ^17^ For example, Philip Morris International replaced flavoured Zyn-branded ONP in California with “flavour ban-approved” Zyn “Smooth” and “Chill” products. Zyn Chill contains a synthetic cooling agent, WS-3, that lacks menthol’s minty odor, but retains its cooling chemesthetic properties.^17^ While California legislators subsequently banned all cooling characterizing flavours, the industry continues to market such ONP, switching to concept flavour names (e.g. from Zyn “Chill” to Zyn “Classic”) that obfuscate the presence of flavour chemicals.^18^ ^19^

With cooling chemesthetic flavourants challenged by regulations, tobacco companies may seek out flavourants with other chemesthetic properties to increase the appeal of ONP and bypass flavour bans. In a web search for newly introduced US-marketed “unflavored” tobacco products, we came across a new ONP product, Lucy “Heat”, that exemplifies such a novel strategy. We investigate the marketing of the product, test its functional effects on chemesthetic receptors that elicit heat sensation and determine the presence of chemesthetic flavourants activating such receptors by chemical analysis, comparing to other newly introduced “spicy” or “heat”-flavoured ONP marketed with additional characterizing flavours.

## METHODS

### ONP product purchases

Lucy Heat and Clear pouches were ordered on the Lucy brand website (lucy.co) in April 2026 and shipped by the company Nicobolt.com. Velo Plus Tropical Heat, Zone Jalapeño Lime, Spicy Mango and Spicy Strawberry pouches were purchased in June and July 2026 from Nicokick.com. Velo Plus Smooth and Zone White pouches were purchased at a convenience store in San Francisco, California, in June 2026.

### Analysis of chemesthetic receptor sensitivity to ONP extracts by calcium microfluorimetry

ONP contents were stirred overnight in 4 mL calcium (Ca^2+^) assay buffer (Hank’s Balanced Salt Solution with 10 mM HEPES (N-2-hydroxyethylpiperazine-N-2-ethane sulfonic acid)) and dilutions of these extracts (diluted 20X-2,000,000X in assay buffer) were prepared to test for receptor activity. The 20X dilution is defined as the extract of one ONP pouch in 20 mL assay buffer, and 2000X the 100-fold dilution thereof. Chemesthetic activity of ONP extracts (diluted 20X-2,000,000X) was tested by intracellular calcium microfluorimetry in HEK-293t cells (RRID:CVCL1926) expressing the human heat/pain receptor, TRPV1 (hTRPV1), as described earlier.^20^ ^21^ Briefly, hTRPV1 expressing HEK-293t cells plated in 96-well plates and loaded with fluorescent Ca^2+^ indicator dye, Calcium 6 (Molecular Devices), were exposed to various dilutions of ONP extracts and specific agonists. Ca^2+^ influx into the cells was monitored in a fluorescent plate reader FlexStation III (Molecular Devices). To control for any variations in receptor expression levels and loading of Ca^2+^indicator across experiments, Ca^2+^-influx data was normalised to the Ca^2+^-response elicited by a maximally activating concentration of agonists (Capsaicin (3 µM)). Responses from cells transfected with empty vector (pcDNA3.1) were recorded as negative controls. Specificity for TRPV1 was validated by use of the selective TRPV1 inhibitor, SB366791 (Tocris Bioscience).^22^

Dose-response curves for receptor activity and associated calcium influx changes were plotted using non-linear regression analysis with a 3-or 4-parameter logistic equation (GraphPad Prism V.11.0, San Diego, California, USA). To further characterise receptor activation by ONP extracts, we determined and compared their efficacies (maximal receptor activation) and potencies (the concentration that produces 50% of its maximal response at a receptor) of activation. Experiments were repeated no less than three times (n ≥ 3) and each conducted at least in duplicates or triplicates (N ≥ 2) with independent ONP extractions.

### Quantification of capsaicinoids in ONP

Capsaicinoid standards were Capsaicin (Sigma), Dihydrocapsaicin (Cayman Chemical Company), Nordihydrocapsaicin (Cayman) and Nonivamide (TCI America) purchased at ≥98% purity. ONP contents were weighed and extracted in 10 mL of methanol for 8 hours to overnight in the dark with constant shaking using a low-speed shaker. Extracts were filtered using a 0.45 µm PVDF filter. Samples were analysed by Ultra-Performance Liquid Chromatography – Mass Spectrometry – Mass Spectrometry (UPLC-MS/MS) by injecting 1µl of diluted samples into a Waters Acquity I-class plus UPLC with a Waters Acquity UPLC BEH C18 column with mobile phase A (0.1% formic acid in water) and mobile phase B (0.1% formic acid in acetonitrile) at 50°C column temperature. Multiple Reaction Monitoring (MRM) mass spectra were acquired with Sciex Triple 7500+ mass spectrometer (Framingham, MA) under positive mode electrospray ionization, with Sciex OS 4.0.2.188 software used for data acquisition. Data analysis was performed using Skyline software (www.skyline.ms). Recovery of capsaicinoids was 100.1±2.8 and 99.0±1.9 % for Lucy and Zone ONP, respectively, when known amounts of nonivamide (10 µg) and capsaicin (10 µg) were spiked into their corresponding ONP extract matrices (∼20 µg/pouch was spiked with a recovery of ∼20.02±0.6 µg/pouch for Lucy ONP and ∼19.8±0.4 µg/pouch for Zone ONP). Limit of quantification (LOQ) for nonivamide analysis in Lucy Heat pouches was 50 ng/mL. LOQ for nonivamide for other pouches analyzed was 0.5 ng/mL. LOQ for all other capsaicinoids tested was 0.5 ng/mL. Samples were analysed from 2-4 independent extractions.

### Quantification of artificial sweeteners in ONP

ONP product extracts were prepared by stirring the contents of one pouch overnight at room temperature in 10 mL of ultrapure water (Thermo Fisher Scientific). Extracts were centrifuged and supernatant and collected for simultaneous quantification of two high-intensity synthetic sweeteners, acesulfame-k (Sigma Aldrich LLC., Saint Louis, MO; ≥99% purity) and sucralose (Sigma; ≥98% purity) by a modified Liquid Chromatography - Mass Spectrometry (LC-MS) methodology used previously. ^23^ ^24^

### Determination of spiciness of ONP on the Scoville scale

Scoville Heat Units (SHU) for each capsaicinoid-containing pouch were calculated from UPLC-determined capsaicinoid concentrations using a modified weight-based approach adapted from the American Spice Trade Association (ASTA) Method 21.3 for assessing the pungency of capsicum products.^25^ The standard ASTA calculation, which incorporates the major naturally occurring capsaicinoids capsaicin, dihydrocapsaicin, and nordihydrocapsaicin, was adapted to include the synthetic capsaicinoid nonivamide. Concentrations of capsaicin, dihydrocapsaicin, nordihydrocapsaicin, and nonivamide were converted to SHU using their respective pungency factors (16, 16, 9.3, and 9.2 SHU per ppm) and summed to estimate total product pungency.^25^ ^26^ Total SHU was calculated as:

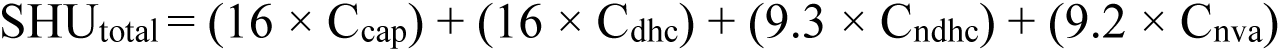

where C_cap_, C_dhc_, C_ndhc_ and C_nva_ represent the concentrations (µg/g pouch; ppm) of capsaicin, dihydrocapsaicin, nordihydrocapsaicin, and nonivamide, respectively. Capsaicinoid quantities measured on a per-pouch basis were normalized to pouch mass (µg/g) prior to SHU calculation. The resulting SHU values were categorized as non-pungent (0-700 SHU), mildly pungent (700-3,000 SHU), moderately pungent (3,000-25,000 SHU), or highly pungent (25,000-70,000 SHU).^27^ ^28^ SHU ratings of ONP products were compared with published SHU values determined for jalapeño chili peppers, a widely consumed pungent food product. Jalapeño SHU values were based on the concentration of capsaicinoids measured per gram of pepper dry weight.^29^

## RESULTS

### Identification of “unflavoured” Lucy Heat pouches sold exclusively in California, Massachusetts and the District of Columbia

In a search of major tobacco product brand websites in May 2026 for products marketed as “unflavoured” we noticed a new link on the website for Lucy products (Lucy.co), a major brand selling ONP, flavoured nicotine gum and lozenges popular among US adolescents and young adults.^30–32^ The link is labelled “California, Massachusetts or DC resident? Try LUCY Unflavoured Product” (figure 1A).^33^. This link directs customers to a page listing three “unflavoured” ONP varieties, “Tobacco”, “Clear” and “Heat”, in nicotine strengths of 4, 8 and 12mg, indicating “Currently available to California, Massachusetts, and Washington, D.C. customers.” (figure 1B).^34^ While products with “Tobacco” or “Clear” labels, indicating tobacco flavour or no flavour, are widely marketed in states banning characterizing flavours, this was the first time we encountered an “unflavoured” ONP labelled exclusively with the descriptor “Heat”. ^34^

**Figure 1.**
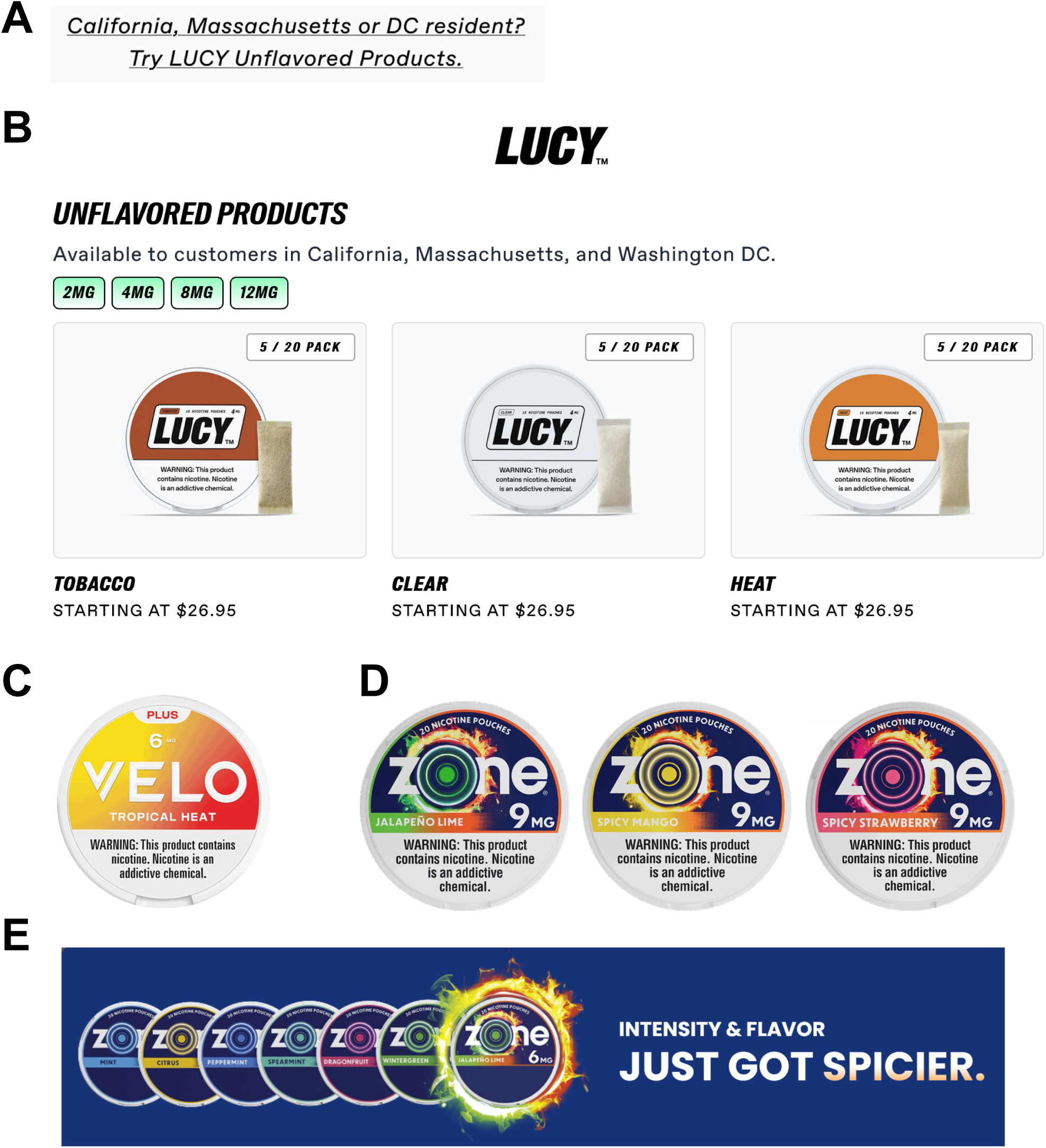
Advertising claims for Heat and Spicy Oral Nicotine Pouches. **A.** Link text on Lucy brand website guiding users to “unflavoured” products sold to California, Massachusetts and District of Columbia customers. **B.** Lucy brand sales site listing “unflavoured” ONP, including “tobacco”, “clear”, and the new “heat” variety. **C.** Image of can of Velo “Tropical Heat” pouches, retrieved from Nicokick.com vendor website. **D.** Images of “Spicy” Zone ONP varieties, “Jalapeño Lime”, “Spicy Mango” and “Spicy Strawberry”, retrieved from Nicokick.com vendor website. **E.** Advertising image from Zone ONP website introducing Zone “Jalapeño Lime” with “Intensity & Flavour Just Got Spicier.”

Searches of ONP vendor websites with the terms “Heat” and related terms “Hot” or “Spicy” revealed additional recently introduced ONP varieties. However, these were advertised with additional concept or characterizing flavours. These products include Reynolds American’s Velo Plus Tropical Heat, introduced in late 2025 and advertised featuring “A bold, juicy mango flavour with just a hint of spice” and Imperial Tobacco Group’s Zone brand introduced Jalapeño Lime, Spicy Mango and Spicy Strawberry varieties in 2026.^35^ ^36^ Zone Jalapeño Lime was announced on the Zone home page with “Intensity and flavour just hot spicier” and is heavily advertised through NASCAR team sponsorship.^36^ ^37^ Zone Spicy Mango and Zone Spicy Strawberry are exclusively sold through the online merchant, Nicokick, a major vendor of oral nicotine products in the US.^36^ ^38^ The vendor announces that “Spicy nicotine pouches are one of the newest flavour categories available at Nicokick. Alongside familiar options like mint, fruit, citrus, coffee, and cinnamon, this category features tobacco leaf-free nicotine pouches with heat-inspired flavour profiles. Intensity and flavour just got spicier”.^38^

### Effects of Heat / Spicy ONP extracts on the chemesthetic receptor TRPV1

The chemesthetic flavourants that elicit the sensations of heat and spicy pungency are the capsaicinoids, a group of natural products generated by chili pepper plants. Capsaicinoids activate Transient Receptor Potential Vanilloid 1 (TRPV1), a heat-sensitive calcium-permeable ion channel expressed in nociceptors, the sensory nerves signaling pain.^39^ ^40^. To examine whether capsaicinoids are present in the Heat and Spicy ONP varieties, serial dilutions of aqueous extracts from Lucy Heat, Velo Plus Tropical Heat and Zone Jalapeño Lime were tested by Ca^2+^ microfluorimetry of HEK293t cells expressing the human TRPV1 receptor. Lucy Heat extract robustly activated TRPV1, with significant activation even at 1:20,000 dilution (equivalent to extract of 1 ONP in 20 liters buffer) (figure 2A). The maximal efficacy of Lucy Heat ONP extracts at TRPV1 was similar to a maximally activating capsaicin concentration (3 µM; ∼103±4%, 95% CI: 94-112%). Control cells transfected with empty expression vector did not display any Ca^2+^-influx in the presence of Lucy Heat extracts (figure 2A). TRPV1 activation by Lucy Heat ONP extracts was significantly attenuated in the presence of the competitive TRPV1 antagonist, SB266791, demonstrating that Ca^2+^-influx activated by Lucy Heat extracts is mediated by TRPV1 (figure 2A). Extracts from Velo Plus Tropical Heat also activated TRPV1, but with less potency than Lucy Heat extracts, with increase in activity observed at significantly lower dilutions than for Lucy Heat extracts, at 1:200 (1 ONP in 200 mL) (figure 2B). Extracts from Zone brand ONP Jalapeño Lime, Spicy Mango and Spicy Strawberry also activated TRPV1, with less potency than Lucy Heat and higher potency than Velo Plus Tropical Heat, with dilutions of 1:600 and 1:2000 (1 ONP in 600 or 2,000 mL) producing significant activity (figure 2C). SB266791 also abrogated these responses, demonstrating that the activation is specifically mediated by TRPV1 (figure 2B,C). Control extracts from other ONP marketed as “unflavoured” with non-heat flavour descriptors, Velo Plus Smooth (figure 2B), Lucy Clear, and Zyn Smooth produced no specific TRPV1 activity (data not shown). Interestingly, Zone White ONP, also marketed as “unflavoured” but without heat flavour descriptors, elicited robust TRPV1-mediated Ca^2+^ influx (figure 2D). Lucy Heat extracts were also the most efficacious (∼103% TRPV1 activation at 1:200 dilution vs 3 µM capsaicin), followed by Zone products (∼65-75%) and Velo Plus Tropical Heat (∼40%). Overall, these results suggest that “Heat” and “Spicy” ONP, along with Zone White, contain hot capsaicinoids.

**Figure 2.**
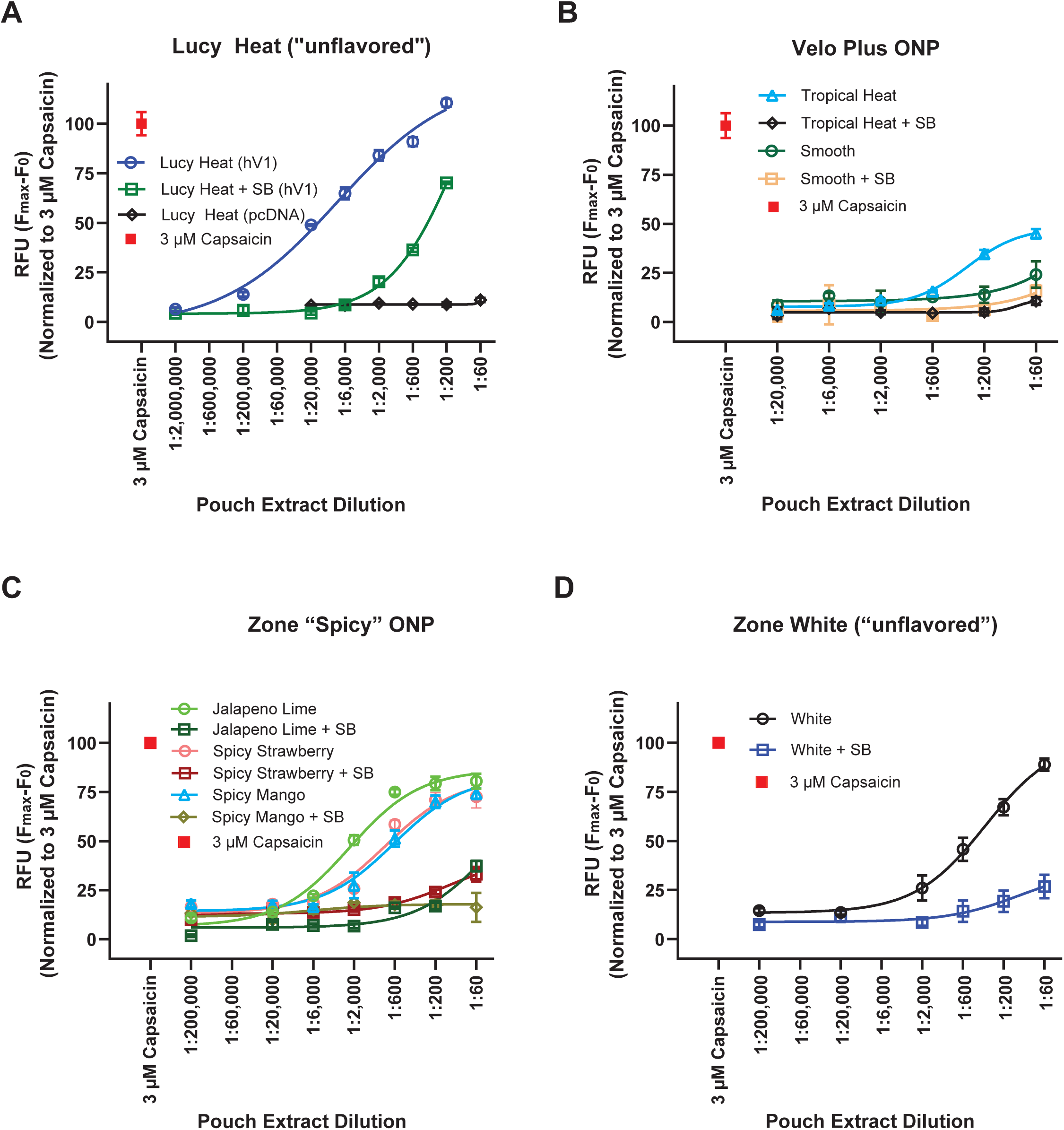
Chemesthetic activity of “Heat” and “Spicy” ONP measured by Ca^2+^ microfluorimetry in HEK293 cells. **A.** Dose-response analysis of human TRPV1 heat/pain receptor-mediated Ca^2+^-influx into HEK293t cells following superfusion with a dilution series of “unflavoured” Lucy Heat ONP extracts (1:200 to 1:2,000,000) in the absence (blue) and presence (green) of competitive TRPV1 antagonist, SB366791 (10 µM). Responses from cells transfected with empty vector (pcDNA3.1) were recorded as negative controls (black). The increase in fluorescence units (F_max_-F_0_) of the fluorescent Ca^2+^ indicator (Calcium 6), was normalised to the Ca^2+^-response elicited by a saturating concentration of agonist capsaicin (3 µM, red). **B.** Dose-response analysis of human TRPV1 heat/pain receptor-mediated Ca^2+^-influx into HEK293t cells following superfusion with a dilution series (1:60 to 1:20,000) of Velo Plus Tropical Heat ONP and Velo Plus Smooth in the absence (Tropical Heat in light blue; Smooth in green) and presence (Tropical Heat in black; Smooth in light orange) of competitive TRPV1 antagonist, SB366791 (10 µM), normalised to the Ca^2+^-response elicited by a saturating concentration of agonist capsaicin (3 µM, red). **C.** Dose-response analysis of human TRPV1 heat/pain receptor-mediated Ca^2+^-influx into HEK293t cells following superfusion with a dilution series (1:60 to 1:200,000) of Zone Jalapeño Lime, Spicy Strawberry or Spicy Mango in the absence and presence of competitive TRPV1 antagonist, SB366791 (10 µM), normalised to the Ca^2+^-response elicited by a saturating concentration of agonist capsaicin (3 µM, red). **D.** Dose-response analysis of human TRPV1 heat/pain receptor-mediated Ca^2+^-influx into HEK293t cells following superfusion with a dilution series (1:60 to 1:200,000) of “unflavoured” Zone White ONP in the absence and presence of competitive TRPV1 antagonist, SB366791 (10 µM), normalised to the Ca^2+^-response elicited by a saturating concentration of agonist capsaicin (3 µM, red). Dose response experiments for TRPV1 activation were performed 2-4 times in duplicates or triplicates with independent ONP extractions, and for experiments in the presence of antagonist SB366791 were performed 1-4 times in duplicates or triplicates. Error bars for each data point represent standard error of the mean. Graphs shown here are from a representative experiment.

### Capsaicinoids in ‘Heat’ and ‘Spicy’ ONP

The major hot capsaicinoids present in chili peppers are capsaicin (CAP), dihydrocapsaicin (DHC) and nordihydrocapsaicin (norDHC).^41^ Another capsaicinoid that is frequently used as a food additive is nonivamide (NVA), often produced synthetically. ^42^ Heat or ‘hot’ and ‘spicy’ ONP were analyzed for these flavourants by UPLC-MS/MS. Analysis of the Lucy Heat ONP demonstrated the presence of nonivamide (675.7±55 μg/pouch) as the only major capsaicinoid (figure 3A). Lucy Heat contained only trace amounts (50-150 ng/pouch) of other capsaicinoids screened for, amounts several thousand-fold lower than nonivamide (figure 3B). Nonivamide and other capsaicinoid contents in Lucy Heat ONP did not differ across nicotine strengths (figure 3A and 3B and table 1). Capsaicinoids were also detected in Zone Jalapeño Lime (NVA: 9511±664; CAPS: 8490±898; DHC: 4010±320; and norDHC: 408±30 ng/pouch), Zone Spicy Mango (NVA: 11821±541 ng/pouch) and Spicy Strawberry (NVA: 15970±1260 ng/pouch) (figure 3D; table 1), with similar contents across nicotine strengths (figure 3D; table 1). Similarly, capsaicinoids were also detected in Velo Plus Tropical Heat (NVA: 116±5; CAPS: 2869±161; DHC: 2200±102; and norDHC: 391±222 ng/pouch) (figure 3C; table 1). Overall, total capsaicinoid contents in Zone ONPs and Velo Plus Tropical Heat were much lower than in Lucy Heat.

**Figure 3.**
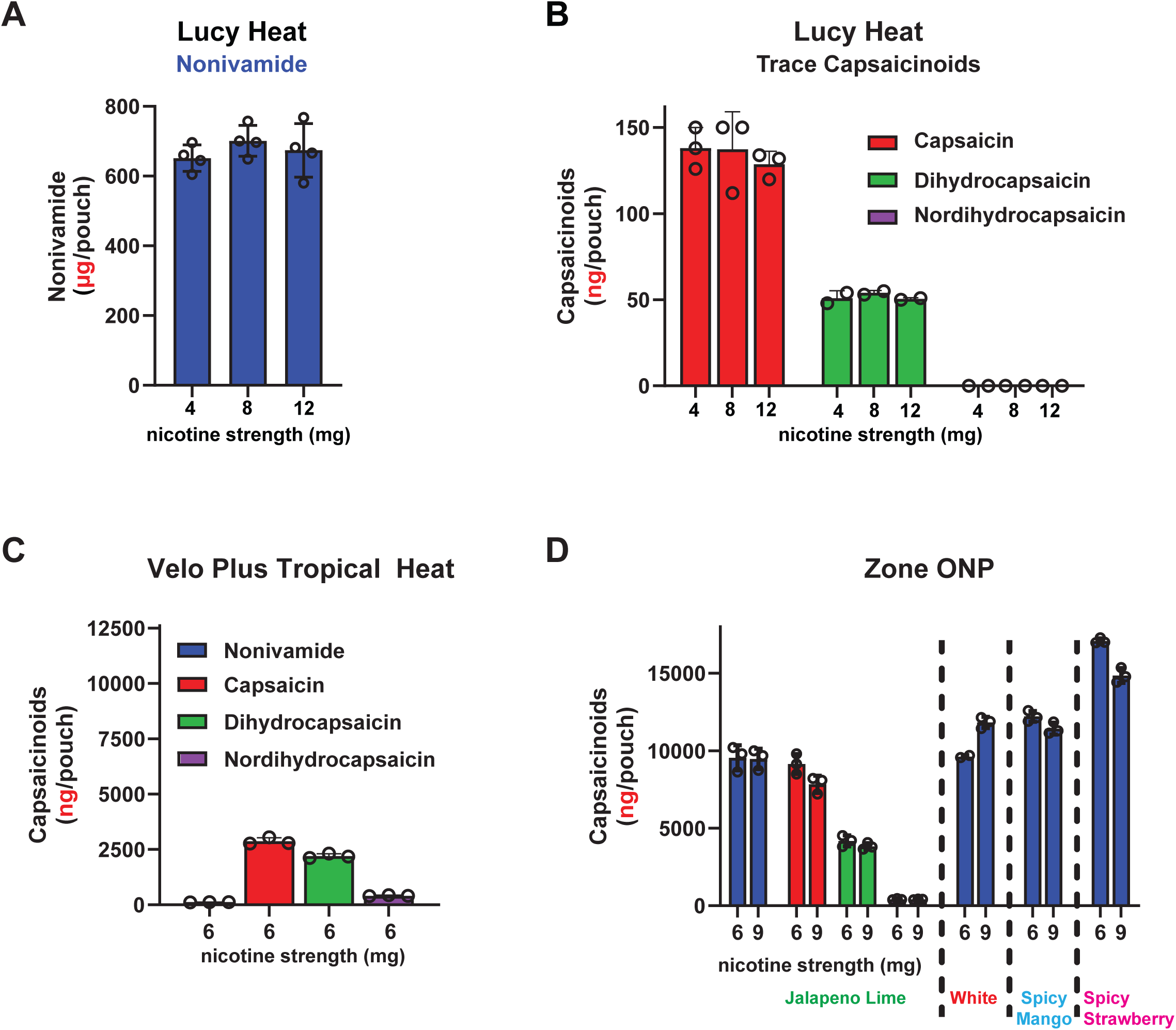
Capsaicinoid contents in “Heat” and “Spicy” ONP measured by UPLC-MS/MS. **A.** Nonivamide contents in Lucy “Heat” ONP at 4mg, 8mg and 12mg nicotine strengths, displayed as microgram (µg) per pouch. **B.** Trace capsaicinoids capsaicin (red), dihydrocapsaicin (green) and nordihydrocapsaicin contents in Lucy “Heat” ONP at 4mg, 8mg and 12mg nicotine strengths, displayed as nanogram (ng) per pouch. **C.** Capsaicinoids in Velo Plus “Tropical Heat” pouches (6 mg nicotine strength), displayed in nanogram (ng) per pouch. **D.** Capsaicinoids in Zone “Jalapeño Lime”, “White”, “Spicy Mango” and “Spicy Strawberry” pouches (6 mg and 9 mg nicotine strengths), displayed in nanogram (ng) per pouch. Chemical analysis was performed from 2-4 independent ONP extractions. Error bars represent standard deviation of the mean.

**Table 1:** Capsaicinoid contents and pungency of US-marketed ONP with ‘Heat’ or ‘Spicy’ flavour descriptors, along with reference products without heat descriptors. Amounts listed for each chemical are per pouch, along with ± standard deviation (n=3-4). <LOQ is below level of quantification. Pungency is listed in Scoville heat units (SHU).

| **Brand** | **Flavor Descriptor** | **Nicotine strength (mg)** | **(ng/Pouch; ± SD)** |  |  |  | **Scoville Heat Units (Pungency)** |
|---|---|---|---|---|---|---|---|
|  |  |  | **Nonivamide** | **Capsaicin** | **Dihydro-capsaicin** | **Nordihydro-capsaicin** |  |
| Lucy | Heat (Unflavored) | 4 | 651688 ±37847 | 138.0±12 | 51.0±4 | <LOQ | 10,712 (**moderate**) |
|  |  | 8 | 701250 ±44456 | 137.3±22 | 54.0±1 | <LOQ | 11,666 (**moderate**) |
|  |  | 12 | 674175 ±76940 | 128.7±8 | 50.5±1 | <LOQ | 11,215 (**moderate**) |
|  | Clear (Unflavored) | 4 | <LOQ | <LOQ | <LOQ | <LOQ | 0 |
| Velo Plus | Smooth (Unflavored) | 6 | <LOQ | <LOQ | <LOQ | <LOQ | 0 |
|  | Tropical Heat | 6 | 116±5 | 2869 ±161 | 2200±102 | 409.0±25 | 180 (non-pungent) |
| Zone | Tobacco | 6 | <LOQ | <LOQ | <LOQ | <LOQ | 0 |
|  | White (Unflavored) | 6 | 9629±87 | <LOQ | <LOQ | <LOQ | 221 (non-pungent) |
|  |  | 9 | 11825±364 | <LOQ | <LOQ | <LOQ | 270 (non-pungent) |
|  | Jalapeno Lime | 6 | 9549±815 | 9145±649 | 4182+351 | 409.3±38 | 750 (mild) |
|  |  | 9 | 9473±658 | 7834±552 | 3838±211 | 407.3±29 | 694 (non-pungent) |
|  | Spicy Mango | 6 | 12215±346 | <LOQ | <LOQ | <LOQ | 293 (non-pungent) |
|  |  | 9 | 11426±380 | <LOQ | <LOQ | <LOQ | 282 (non-pungent) |
|  | Spicy Strawberry | 6 | 17083±171 | <LOQ | <LOQ | <LOQ | 390 (non-pungent) |
|  |  | 9 | 14857±474 | <LOQ | <LOQ | <LOQ | 342 (non-pungent) |
| Zyn | Smooth (Unflavored) | 3 | <LOQ | <LOQ | <LOQ | <LOQ | 0 |
|  | Original (Unflavored) | 3 | <LOQ | <LOQ | <LOQ | <LOQ | 0 |
| Jalapeno Pepper (Reference)# |  |  | 17,400 ±17,500 | 800,000 ±590,000 | 547,000 ±406,000 | 101,000 ±82,100 | 21,700 ± 15,500 (**moderate**) |

<sup>#</sup> Capsaicinoid contents and pungency levels for jalapeño pepper (per gram of pepper dry weight) were derived from Sweat et al., 2016. <sup>29</sup>

No capsaicinoids were detected in Zyn Smooth with our method, whereas their levels were close to the limit of quantification (LOQ) in Velo Plus Smooth. Interestingly, significant amounts of NVA (10946±1231 ng/pouch; table 1) were detected in Zone White, the Zone ONP variant marketed as “unflavoured” without heat descriptors, validating calcium microfluorimetry data (figure 2D).

### Spiciness of Heat”, “Spicy” and “Unflavoured” ONP on the Scoville scale

The mean spiciness of Lucy Heat products on the Scoville scale was determined as 11,197 ± 477 SHU across nicotine strengths, placing them in the moderately pungent category (3,000-25,000 SHU) and within the range reported for jalapeño chili peppers (21,700 ± 15,500 SHU). ^29^ In contrast, the average pungency of Zone Jalapeño Lime products across nicotine strengths was 722 ± 39 SHU, which would be classified as mildly pungent. Among the remaining tested products in which capsaicinoids were detected, including Velo Plus Tropical Heat, Zone’s Spicy Mango, Spicy Strawberry, and White, pungency ranged from 180 to 390 SHU, indicating a warming effect, but little to no perceptible pungency.

### Artificial Sweeteners in “Heat”, “Spicy” and “Unflavoured” ONP

The artificial sweeteners acesulfame K (AceK) and sucralose are widely used in ONP of all major brands, and AceK is a declared ingredient in Lucy Heat ONP. ^23^ ^43^ AceK and sucralose were quantified in both “unflavoured” Lucy Heat and Zone White, and in Zone Jalapeño Lime ONP. AceK, but not sucralose, was confirmed to be present in Lucy Heat ONP across all nicotine strengths (∼2.1-3.3 mg/pouch), whereas sucralose, but not AceK, was detected in Zone White and Zone Jalapeño Lime ONP, between ∼1.7-2.1 mg/pouch (supplementary table 1).

## DISCUSSION

This study describes a new flavour category of ONP, “Heat” or “Spicy” pouches, introduced in late 2025 and in 2026. We demonstrate that these products contain hot and pungent capsaicinoids, the flavourants produced by chili peppers, that activate the heat-and pain receptor, TRPV1, in peripheral sensory neurons. Two of these products, Lucy Heat and Zone White, marketed as “unflavoured”, are specifically targeted towards consumers in United States jurisdictions that implemented bans on flavoured tobacco products. In other products, such as Velo Plus Tropical Heat and Zone Jalapeño Lime, capsaicinoids are combined with additional characterizing flavours (mango, lime).

Lucy Heat almost exclusively contains nonivamide as hot and pungent flavourant, a synthetically produced capsaicinoid authorized for use as a food additive, also present in pepper spray and related self-defense or riot control weapons.^42^ ^44–46^ The average nonivamide content of Lucy Heat ONP was 675.7 μg/pouch across all nicotine strengths. Only trace amounts of other capsaicinoids were detectable that are likely impurities from synthesis.^42^ Lucy Heat contained more than 33 times more capsaicinoids than the next-most potent product tested, Zone Jalapeño Lime, at 22.4 μg/pouch total capsaicinoids, 57 times more nonivamide than Zone White at 11.8 μg/pouch and 121 times more capsaicinoids than Velo Plus Tropical Heat, at 5.5 μg/pouch total capsaicinoids. In addition to nonivamide, Zone Jalapeño Lime and Velo Plus Tropical Heat ONP contain significant amounts of capsaicin, dihydrocapsaicin and nor-dihydrocapsaicin, suggesting these products may contain natural chili pepper extracts or spices, possibly in combination with synthetic nonivamide.

Extracts from Lucy Heat products robustly activated human TRPV1 receptors, even at a dilution of 1:20,000. The human sensory detection threshold for capsaicin in the mouth was determined to be around 0.9 µM (micromolar).^47^ Nonivamide is 1.72-fold less potent than capsaicin at human TRPV1 receptors, suggesting a higher oral detection threshold for nonivamide of approximately 1.5 µM.^48^ In humans, resting saliva production (RSP) was determined to be 0.6 ± 1.2 mL in 5 min, while stimulated saliva production (SSP) was 8.5 ± 4.6 mL/5 min.^49^ Given that ONP use and chemesthetic flavourants activate saliva flow, and assuming that all of the 675.7 μg/pouch nonivamide is released from the Lucy Heat pouch, the concentration of nonivamide in stimulated saliva production would be approximately 270 µM, exceeding its detection threshold by 180-fold.^50^ ^51^ Nonivamide may be released more slowly, with the highest concentration at the pouch insertion site between gum and lip. Based on these estimates, Lucy Heat pouches likely impart a noticeable hot and pungent flavour.

To further classify the pungency of the tested ONP products, we converted their capsaicinoid contents into Scoville Heat Units (SHU), the units on the Scoville scale of spiciness used by growers and spice manufacturers that ranks chili peppers and pepper-containing products according to their spiciness. ^29^ ^52^ Lucy Heat ranks within the “moderately pungent” category, similar to dried Jalapeño peppers. Based on this analysis, we think that Lucy Heat products might appeal to users who tolerate spicy foods, but not to people who avoid noticeable spiciness. All other products tested (Velo and Zone ONP) ranked one category below (mildly pungent), and, besides “unflavored” Zone White, contain fruit (mango, strawberry) and citrus (lime) flavour notes that likely have broader appeal, especially for youth who prefer fruit flavoured ONP.^4^ ^8^ Recent United States National Youth Tobacco Survey (NYTS) data from 2024 and 2025 indicate that ONPs with related “spice” flavours (e.g., cinnamon, vanilla, and clove) are among the most commonly used products by youth, ranking behind only mint/menthol-and fruit-flavored ONP. ^4^ Thus, capsaicinoid-flavoured ONP may strongly appeal to youth who often experiment with new and unusual flavour combinations. Human studies evaluating the sensory and perceptual attributes of these products, especially among youth and young adults, would provide valuable insight into their consumer appeal and use experience.

Artificial sweeteners were detected in significant quantities in the ONP tested in this study. While Lucy Heat ONP contained AceK, Zone Jalapeño Lime and White contained sucralose. These artificial sweeteners are widely used in ONP of the major US-marketed brands. ^23^ AceK contents measured in Lucy Heat pouches exceeded the AceK contents (∼≥3-fold) in Zyn ONP with characterising flavours determined by us previously.^23^ The same applies to the sucralose contents in Zone Jalapeño Lime and White that exceeded sucralose contents in previously tested ONP.^23^ Sweet taste is known to partially suppress the burning sensations elicited by oral capsaicin in humans.^53^ ^54^ The manufacturers are likely titrating sweetness vs. irritancy to create a range of products that are palatable for either frequent or initiating users. By combining hot capsaicinoids with sweeteners the manufacturers may also aim to create sensory profiles resembling popular hot and spicy confectionery products, such as Hot Tamales, Atomic Fireballs, and Zotz, which are widely recognized among Generation Z and millennial consumers.

One of the major functions of flavour additives in tobacco products is to mask the adverse effects of nicotine.^13^ Nicotine acts as a sensory irritant creating a burning sensation in the mouth and throat.^55^ The inclusion of capsaicinoid irritants in a tobacco product is a relatively novel product characteristic, and it is unclear how the combination of nicotine and capsaicinoid irritants impacts product appeal. Oral cross-desensitization studies in humans with nicotine and capsaicin indicate a concentration-dependent interaction between both stimuli, with capsaicin reducing nicotine-evoked tongue irritation, whereas only high nicotine concentrations attenuating capsaicin-evoked irritation.^56–58^ These psychophysical studies suggest that nicotine and capsaicinoids do not simply produce additive irritation but rather interact in a complex manner that depends on stimulus intensity and prior exposure.

The introduction of capsaicinoids in ONP also raises toxicological concerns. While capsaicinoids, including the synthetic nonivamide, have received the “Generally Recognized As Safe” (GRAS) designation for use in foods, this safety designation does not apply to tobacco products such as ONP. ^44^ ^59^ Recent studies have revealed that use of ONP can have adverse oral and dental health effects, including gum recession, inflammation, oral microbiome dysbiosis and oral lesions such as leukoplakia, a pre-cancerous lesion that can progress to oral squamous cell carcinoma.^60–66^ Nicotine itself may be a factor in the development of oral leukoplakia.^67^ Pepper extracts and capsaicinoids can induce oral inflammation through neurogenic effects, triggering the release of pro-inflammatory neuropeptides such as Substance P and Calcitonin Gene-Related Peptide (CGRP) from trigeminal nerve endings in the mucosa.^68^ ^69^ Capsaicinoids are also known to affect the proliferation of oral fibroblasts.^70^ While considered safe under their conditions of intended use as flavourants in foods, little is known about how oral mucosa tolerate repeated exposures to capsaicinoids in tobacco products such as ONP, with other sensory irritants such as nicotine and additional flavourants and additives present.^45^ Nicotine is known to sensitise TRPV1 receptors to capsaicinoids, activates trigeminal sensory nerves and triggers nerve-derived CGRP release and may thereby potentiate inflammatory mucosal responses.^55^ ^71–73^ These interactions need to be investigated to minimize the risk this novel category of ONP may pose to public health.

Regulators need to monitor whether capsaicinoids are also added to other tobacco products to produce new flavour varieties. Especially the addition to inhaled products such as electronic cigarettes poses significant risks because of the potent respiratory irritant effects of capsaicinoids.^74^ This is not just a hypothetical concern, as potent irritant flavourants have been added to e-cigarettes before. For example, cinnamaldehyde, a cinnamon flavourant and irritant, was found to be present in certain e-cigarette liquids at high concentrations, causing substantial toxicity.^75^

Lucy Heat is marketed as unflavoured to customers in California, Massachusetts and the District of Columbia. These jurisdictions implemented comprehensive bans of tobacco products with characterizing flavours, with the exemption of tobacco-flavoured products. These bans also extend to the ONP category. Such state and municipal flavour bans have successfully reduced smoking rates and tobacco product sales, including use and sales of modern tobacco products such as electronic cigarettes. ^76–79^ The tobacco industry attempted to bypass flavour bans by introducing chemesthetic flavourants such as synthetic cooling agents replacing menthol and mint flavourants, claiming that chemesthetic flavourants do not impart characterising flavours. ^18^ ^80^ For example, R.J. Reynolds stated that combustible cigarettes containing synthetic cooling flavourants, introduced to replace menthol cigarettes after California’s 2022 flavour ban, do not have a characterising flavour.^14^ California legislators amended legislation to thwart this tactic by clarifying the term “characterizing flavour” to include “a cooling sensation distinguishable by an ordinary consumer during the consumption of a tobacco product.”.^14^ This amendment led to the withdrawal of combustible cigarettes containing cooling flavourants from California, and the rebranding of ONP containing cooling agents with concept flavours.^18^ ^19^

The introduction of “unflavoured” Lucy Heat ONP by the Lucy brand challenges such laws targeting chemesthetic flavourants. The company is apparently convinced that such laws only target cooling flavourants and not capsaicinoids that impart sensations of heat and pungency. Whether such differentiation has legal standing, or if “heat” or “spicy” can stand on their own as a characterising flavour recognized by ordinary consumers, requires further legal and scientific review. Consumers that are widely familiar with the culinary effects of chili peppers and chili pepper-derived flavourants in foods, and, if asked, would likely deem heat and spicyness to be tastes or flavours.

Commercial companies producing flavourants as food additives are using more expansive definitions of the term “flavour”. The *Flavor and Extract Manufacturers Association (FEMA)*, the industry organization responsible for issuing *Generally Recognized as Safe* (GRAS) determinations for food additives in the United States, defines the term “flavour” as “*Flavour is the entire range of sensations that we perceive when we eat a food or drink a beverage. Flavour encompasses a substance’s taste, smell, and any physical traits we perceive in our mouths, such as “heat” (for example, cinnamon) or “cold” (for example, spearmint).*”, clearly including “heat” as a component of the flavour experience.^81^ Major tobacco companies such as Altria, Philip Morris International and R.J. Reynolds are listed as members of FEMA.^82^ It can therefore be assumed that tobacco company personnel determining product flavour compositions are aware, or might have even been involved in the framing of FEMA’s more expansive definition. The FDA was considering a similarly expansive definition of the term “characterizing flavour” in its proposed Standard for Menthol Cigarettes, including “*The presence and amount of artificial or natural flavour additives…; The multisensory experience (i.e., taste, aroma, and cooling or burning sensations in the mouth and throat) of a flavour during use of a tobacco product,…;Flavour representations (including descriptors), either explicit or implicit, in or on the labeling (including packaging) or advertising of tobacco products*”, clearly including “burning sensations” as a component of a characterizing flavour.^83^ While the FDA withdrew the rule in January 2025 by authority of the White House Office of Management and Budget (OMB), this proposed rule adds to the evidence for a regulatory and scientific consensus by regulators, legislators and industry that sensations imparted by chemesthetic flavourants, including capsaicinoids, represent characterizing flavours.^84^ In the absence of federal action, US federal states should adopt “characterizing flavor” definitions similar to the language in the proposed federal tobacco product standards for menthol cigarettes to counter industry tactics introducing chemesthetic flavourants to bypass flavour bans.

The presence of sweeteners in “unflavored” capsaicinoid-containing ONP products such as Lucy Heat and Zone White has additional regulatory implications. In FDA’s technical review of the Premarket Tobacco Products Application (PMTA) for Zyn ONP products published in 2025, the technical reviewer stated that “Due to added ingredients such as sweeteners and cooling agents, FDA has determined that all new products have a non-tobacco characterizing flavor for the purposes of this review”.^85^ Thus, based on the presence of sweeteners alone, “unflavoured” Lucy Heat and Zone White ONP would be considered by FDA as having a characterising flavour, if reviewed with the same criteria.

## Funding

This work was supported by grants U54DA036151 (Yale Tobacco Center of Regulatory Science) and grant R01DA060884 from the National Institute on Drug Abuse (NIDA), and grant P30CA014236 (Duke Cancer Institute) from the National Cancer Institute (NCI) of the National Institutes of Health (NIH) of the United States and the Center for Tobacco Products of the US Food and Drug Administration (FDA).

## Disclaimer

The funding organizations had no role in the design and conduct of the study; the collection, management, analysis, and interpretation of the data; the preparation, review, or approval of the manuscript; nor in the decision to submit the manuscript for publication. The content is solely the responsibility of the authors and does not necessarily represent the views of National Institutes of Health or the Food and Drug Administration.

## Competing Interests

S.E.J. receives fees from the California Department of Justice for consulting on matters of tobacco product regulation and serves on the Tobacco Products Scientific Advisory Committee (TPSAC) of the US Food and Drug Administration; he consulted for the World Health Organization and is Vice Chair of the Tobacco Action Committee of the American Thoracic Society (ATS). S.V.J. serves on the Tobacco Action Committee of the American Thoracic Society (ATS).

## Author Contributions

S.V.J. and S.E.J. conceptualized and designed the study. S.E.J. identified and purchased products and wrote the first draft of the paper; S.V.J. provided advice on product choice and carried out product extractions and functional analysis. Z.L. carried out the chemical analysis. S.V. J., Z.L. and S.E.J. contributed to revision of the manuscript. All authors critically reviewed, edited, and approved the final draft before submission. S.E.J. attests that all listed authors meet authorship criteria and that no others meeting the criteria have been omitted.

## Notes

### Summary of Updates

Analytical studies of capsaicinoids in additional Zone ONP products were completed. These data were added to new Figure 3, Panel D. Capsaicinoid contents were converted to Scoville Heat Units on the Scoville Scale of pungency, for easier assessment of product appeal. The discussion was rearranged and revised now including clarification of food safety determinations, discussion of additional toxicological concerns related to electronic cigarettes containing capsaicinoids, and clarification of the effects of artificial sweeteners on product appeal and sensory perception.

