## supplementary table 1 for "Chili Pepper and Synthetic Capsaicinoid Flavourants in “Heat” and “Unflavoured” Oral Nicotine Pouches Marketed in the United States"

**Supplementary Table 1:** Artificial sweeteners in US-marketed Lucy Heat and Zone (White and Jalapeno Lime) ONP. Amounts listed for each chemical are per pouch, along with ± standard error (n=2-3).

| Brand | Flavor Descriptor | Nicotine strength (mg) | mg/pouch (± s.e.) | |
|---|---|---|---|---|
| | | | Acesulfame-K | Sucralose |
| Lucy | Heat | 4 | 3.3±0.8 | |
| | | 8 | 2.7±0.3 | |
| | | 12 | 2.1±0.4 | |
| Zone | White | 9 | | 1.7±0.02 |
| | Jalapeno Lime | 6 | | 2.0±0.04 |
| | | 9 | | 2.1±0.01 |
